# Single cell transcriptional characterization of human megakaryocyte lineage commitment and maturation

**DOI:** 10.1101/2020.02.20.957936

**Authors:** Fizzah A Choudry, Frederik Otzen Bagger, Iain C Macaulay, Samantha Farrow, Frances Burden, Carly Kempster, Harriet McKinney, Lars R Olsen, Ni Huang, Kate Downes, Thierry Voet, Rakesh Uppal, John F Martin, Anthony Mathur, Willem H Ouwehand, Elisa Laurenti, Sarah A Teichmann, Mattia Frontini

## Abstract

In the current understanding of adult bone marrow hematopoiesis, megakaryocytes (MKs) originate from cells immuno-phenotypically indistinguishable from hematopoietic stem cells (HSCs), bypassing intermediate progenitors. Here, we use single cell RNA sequencing to characterize HSCs and MKs from human bone marrow, to investigate MK lineage commitment and maturation. We identify two MK primed HSC clusters exhibiting unique differentiation kinetics, at least one of which is used in steady state and stress thrombopoiesis. By analyzing transcriptional signatures we show that human bone marrow MKs originate from MK primed HSC subpopulations, supporting the notion that these display exclusive priming for MK differentiation. We show that transcriptional programs change with increasing MK ploidy, where genes upregulated in high ploidy states may have functional relevance in platelet production. Finally, we highlight the presence of a specific transcriptional signature in MKs from individuals with myocardial infarction, supporting the aberration of MK differentiation in this thrombotic state.

## Introduction

The classic hierarchical differentiation model of hematopoiesis, in which hematopoietic stem cells (HSC) go through a series of progenitors with increasing lineage restriction^1^ has been challenged with improved cell sorting, functional assays and single cell transcriptome analysis^2–4^. Recent data, based on the transcriptional profiling at single cell level, points to the existence of a hematopoietic stem cell compartment, formed by HSCs and multipotent progenitors, in which gene expression programs driving self-renewal and lineage commitment coexist^5^. Unipotent progenitors differentiate from the stem cell compartment and further lineage commitment is marked by key transcripts^2–3^. One of the first evidence against the classical model of hematopoiesis was the identification of lymphoid primed multipotent progenitor (LMPP) in mouse^5,6^, and multi lymphoid progenitor (MLP) in human^7^, both showing an early loss of megakaryocyte-erythroid potential. At the same time, megakaryocyte (MK)-primed HSCs were identified in mouse, and shown to transcribe MK/platelet genes, such as VWF^8^ and CD41^9^. More recently, MK unipotent progenitors have also been identified in the phenotypic HSC compartment, where they remain quiescent until activated by acute platelet demand^10^. Furthermore, lineage tracking experiments have established a direct HSC origin, independent from other lineages, for at least half of the MKs^11^. These findings complement early evidence of a direct functional relationship between MK and HSC; with MKs inhibiting HSC proliferation via thrombopoietin (TPO)^12,13^ and CXCL4 (PF4)^14,15^, as well as, their transcriptional similarity^16^, dependence on shared transcription factors^17^ and proximity within the stem cell niche^18^.

The primary physiological function of MKs is thrombopoiesis where each cell releases up to 6,000 platelets into the bloodstream. MKs comprise <0.01% of the total number of nucleated cells in the bone marrow and are the largest cell residing in the bone marrow with diameters up to 150μm. They undergo multiple rounds of DNA replication without cell division, known as endomitosis, resulting in cells with an enlarged cytoplasm and average ploidy, in man, of 16N^19^. The complex MK-platelet hemostatic system is unique to mammals^20^ and the function of MK polyploidization and its evolutionary advantage remains unclear. A number of technical challenges have precluded the study of human bone marrow residing MKs including recruitment of healthy bone marrow donors, the rarity of the cell population, and their size and fragility. These have so far prevented transcriptome analysis of primary human bone marrow MKs. Much of the current knowledge on MKs is based on gene expression array data of *in vitro* differentiated MKs cultured from CD34+ cells obtained from fetal liver, cord blood and adult blood, despite the fact that they are known to be phenotypically distinct from their *in vivo* counterparts^21–25^. In contrast to human bone marrow MKs, *in vitro* differentiated MKs cultured from CD34+ cells have average ploidy levels of 2N and limited capacity for platelet production^26^. It is therefore likely that there are drivers of differentiation, ploidy increase, and maturation of *in vivo* MKs that have yet to be identified.

Platelets play a pivotal role in hemostasis by surveying the vasculature for endothelial lesions and tissue inflammation. When activated they adhere, spread and form thrombus, in order to maintain vascular integrity. Aberration in platelet count and function can result in thrombotic or bleeding disorders^27^. During steady state thrombopoiesis, a constant circulating platelet mass is maintained by an inverse relationship between platelet count and volume. There is a complex homeostatic control system which is in part, but not solely, mediated by TPO where a change in hemostatic demand leads to modulation of TPO levels which regulate MK maturation^28^ and HSC quiescence^14^. This results in the acute release of platelets of greater volume to maintain overall circulating platelet mass^27^. Evidence suggest that stem-like megakaryocyte committed progenitors may be activated upon acute inflammatory stress^10^. Myocardial infarction has also been suggested as a model for stress thrombopoiesis based on the observation of elevated mean platelet volume and reticulated/immature platelet fraction in the acute setting suggesting an upregulation of the HSC-MK axis^29^. However, it is not clear whether “emergency” or “stress” thrombopoiesis is due to feedback at the level of HSC differentiation, or at the MK level.

Here, we performed single cell RNA sequencing on human bone marrow HSCs and MKs and found that the HSC population could be divided into 5 clusters, based on their gene expression signatures, and that two of the clusters represent MK primed HSC. Human bone marrow MKs were also sequenced, single cell and low input, and were shown to branch from these MK primed HSC clusters using differentiation trajectory analysis. Primary MK transcriptome analysis demonstrated that, with increasing ploidy, there are two distinct transcriptional states. Increasing MK ploidy is associated with downregulation of platelet specific genetic programs and the upregulation of translation and protein localisation, as well as, with the expression of a number of transmembrane receptors implicated in platelet production. Finally, we analyzed the primary MK transcriptome in the clinical setting of myocardial infarction and found changes in gene expression landscape, supporting the notion of a role for stress thrombopoiesis in this pathological state.

## Results

### Single cell RNA sequencing of *ex vivo* human bone marrow HSCs

To characterize the HSC transcriptional landscape and to investigate early fate commitment events we sequenced 884 single HSCs (CD34+ CD38-CD45RA-CD90+ CD49f+, the compartment most enriched in long-term repopulating HSCs^4^) isolated from fresh bone marrow harvested from 5 individuals undergoing sternotomy for heart valve replacement (Fig. 1a and Supplementary Table S1). The HSCs were index-sorted and the intensities of the signal of cell surface proteins CD34, CD38, CD45RA, CD90 and CD49f were recorded (Supplementary Fig. 1). Raw reads were filtered to exclude cells that gave low quality libraries with a Support Vector Machine (SVM) machine learning method trained on a subset of single cell libraries made from cDNA positive for GAPDH, as measured by qRT-PCR (Supplementary Table S2). This quality filter excluded cells where the majority of the reads mapped to mitochondrial genes or ERCC spike-in controls, both a sign of low cDNA input^30^ while favoring cells that had more reads mapping to exonic regions and to a high number of genes (Fig. 1b), leaving 119 cells that were used for analysis. The SVM selected for larger, more metabolically active cells, while excluding non-viable cells (Fig. 1c). ERCC spike-ins were used to model technical noise^31^ and to identify the 2000 most highly variable genes above the noise model^32^ (Fig. 1d). This filtering strategy slightly favored ribosomal genes. However, analysis of another HSC single cell dataset built using MASS-seq and relying on unique molecular identifier (UMI) count^33^ gave a similar pattern of inter-cellular variance among ribosomal genes (Supplementary Fig. 2).

**Figure 1.**
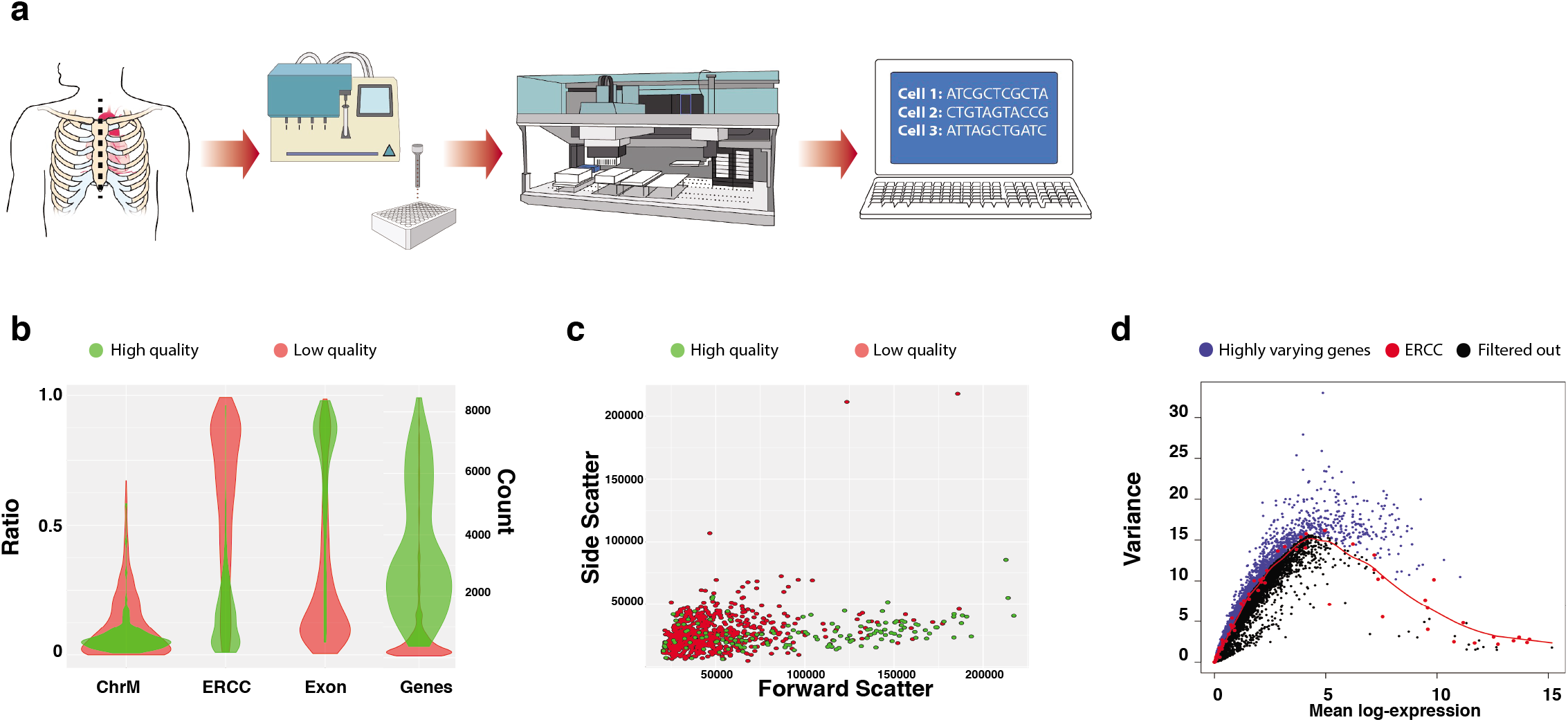
Single cell RNA sequencing of ex vivo human bone marrow HSCs. **a.** Experimental design. Sternal bone marrow was harvested from 5 individuals undergoing sternotomy for heart valve replacement. It was stained with antibodies for surface markers that define the human HSC population and sorted as single cells using index FACS into cell lysis buffer. Single cell RNA sequencing libraries were then prepared, sequenced on a HiSeq 4000 instrument and analysis was performed as described in Methods; **b.** Cell filtering. Performed using a SVM machine learning method (modified from a previously described model^37^) trained on cDNA from HSCs selected by expression of the housekeeping gene *GAPDH.* High and low quality cells shown by mitochondrial mapping (ChrM), ERCC ratio (ERCC), exonic ratio (Exon), number of called genes (Genes). Green denotes high quality, red denotes low quality. Out of 884 cells, 119 were taken forward for further analysis based on high quality; **c.** Filtered cells are larger. Flow cytometric plot FSC/SSC showing cells filtered as high quality by the SVM machine learning method. Green denotes high quality, red denotes low quality; **d.** Gene filtering. Variance of normalized log-expression values for each gene in the HSC dataset, plotted against the mean log-expression. Variance estimates for ERCC spike-in transcripts and curve fit are highlighted in red. Blue dots represent highly varying genes (n=2000).

### Unsupervised clustering of single HSCs: two MK primed HSC subsets

To identify subpopulations of cells within the HSC compartment, we used unsupervised hierarchical clustering of the cells’ Pearson correlation coefficients in the principal component analysis (PCA) space to generate clusters (Fig. 2a) where total silhouette score was used to select the number (from 1 to 20) of clusters tested (Supplementary Fig. 3). We further validated the robustness of our clustering strategy by inspecting the first four principal components (Fig. 2b), where each cluster separated well within the first three components and showed that the first two principal components were not explained by either donor or batch effect (Supplementary Fig. 4). The cells within the HSC population formed 5 distinct clusters. Marker genes for each cluster were then identified based on their predictive value to separate that cluster from the remaining cells (Wilcoxon Rank test p<0.001^8^). The expression of the top 20 genes characterizing each cluster shows distinct expression patterns independently of donor; of note cluster 3 had no significant marker genes (Fig. 2c and Supplementary Table S3 for the full list of genes). Many of the marker genes for cluster 1 and 4 were found to be highly expressed in previously reported MK datasets^25^. Cluster 1 cells were marked by the expression of genes known to be highly expressed in all MK precursors (HSC, multipotent progenitors, common myeloid progenitor, MK-erythroid progenitor) such as *PRKACB, NRIP1, PARP1, HEMGN* as well as *ANGPT1* and *IL1b*, which encode proteins directly involved in platelet function. By contrast cluster 4 cells were characterized only by genes encoding proteins directly involved in platelet function such as: *TUBB1, TUBA4A, F13A1, CCL5* and *GRAP2.*

Gene Ontology terms enrichment analysis of the marker genes revealed early priming patterns: cluster 2 marker genes have an ontological bias towards angiogenesis, endothelial cell function and wound healing, whereas cluster 5 marker genes towards immunity and leukocyte function (Supplementary Tables S4-S7).

### Developmental trajectories of early lineage priming in HSCs

To investigate if the HSCs exhibited lineage priming potential we performed trajectory analysis, independently of the previous clustering, using Monocle2^34^. We obtained potential developmental trajectories where the same clusters re-emerged, with the exception of a few cells (Fig. 2d). This analysis showed two distinct branching points, with clusters 1 and 4 stemming from the same developmental trajectory (Fig. 2d). To determine whether these trajectories could delineate early fate priming in HSC, we derived gene signatures from the highly purified hematopoietic populations in a previously published study^35^ and visualized these signatures onto the trajectory plot. The signature for MK was found enriched in cells localized around branching point 2 and belonging to Cluster 1 and 4 (Fig. 2e), whereas the gene expression signature for the megakaryocyte erythrocyte progenitor (MEP) was found enriched only in cells belonging to cluster 1 after the branching point 2 (Fig. 2f). These signatures are largely non-overlapping (Supplementary Fig. 5) and no single gene appears to drive enrichment seen along the branches (Supplementary Fig. 6). Other hematopoietic signatures overlapped with different clusters, the monocyte gene signature was found enriched in cells belonging to cluster 5 and the lymphocyte signature was found enriched in cells belonging to cluster 2.

### Functional potential of MK primed clusters

We then retrospectively analyzed index fluorescence-activated cell sorting (FACS) data and found that the cells that belong to different clusters based on transcriptome data also show differences based on cell surface markers intensities (Fig. 3a). Cluster 1 and 4 could be identified on the basis of FSC-A and CD34 with cluster 1 having CD34^Lo^ and FSC-A^Hi^, whereas cluster 4 has CD34^Hi^ and FSC-A^Hi^ (Fig. 3b). Analysis of PCA vector loadings showed that the cell surface expression of CD34, FSC-A and CD49f drive most of the variance on PC2/3 within the HSCs (Fig. 3c).

**Figure 2.**
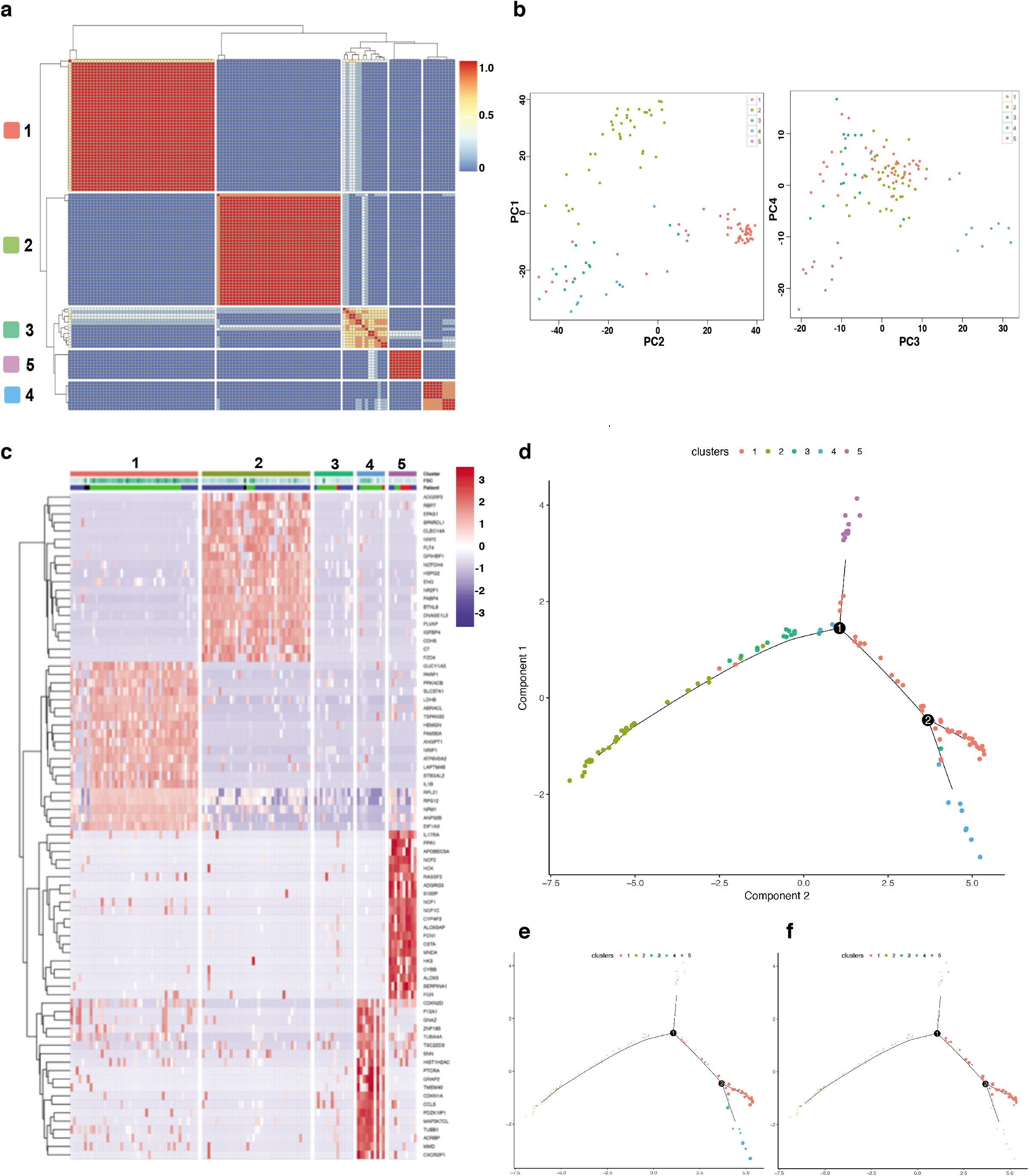
Unsupervised clustering and developmental trajectories of early lineage priming in single HSCs. **a.** Pearson’s correlation map of HSCs in unsupervised clusters by distance within the PCA space, dendrograms are formed by hierarchical clustering on the Euclidean distances between cells; **b.** PCA plots constructed from normalized log-expression values of correlated highly variable genes, where each point represents a cell in the HSC dataset. First, second, third and fourth components are shown; **c.** Heatmap of mean-centered normalized and corrected log-expression values for the top 20 marker genes for each cluster. Dendrograms are formed by hierarchical clustering on the Euclidean distances between genes (row). Column colors represent the cluster to which each cell is assigned; **d.** Ordering single cell differentiation between cell clusters using Monocle 2. Individual cells are connected by a minimum spanning tree with branch points (thin lines), representing the differentiation trajectory and clusters are differentiated by color; **e.** Single cell differentiation trajectory showing expression of MK gene signature from DMAP dataset. Expression of gene signature is indicated by the size of each single cell; **f.** Single cell differentiation trajectory showing expression of MEP gene signature from DMAP dataset. Expression of gene signature is indicated by the size of each single cell.

**Figure 3.**
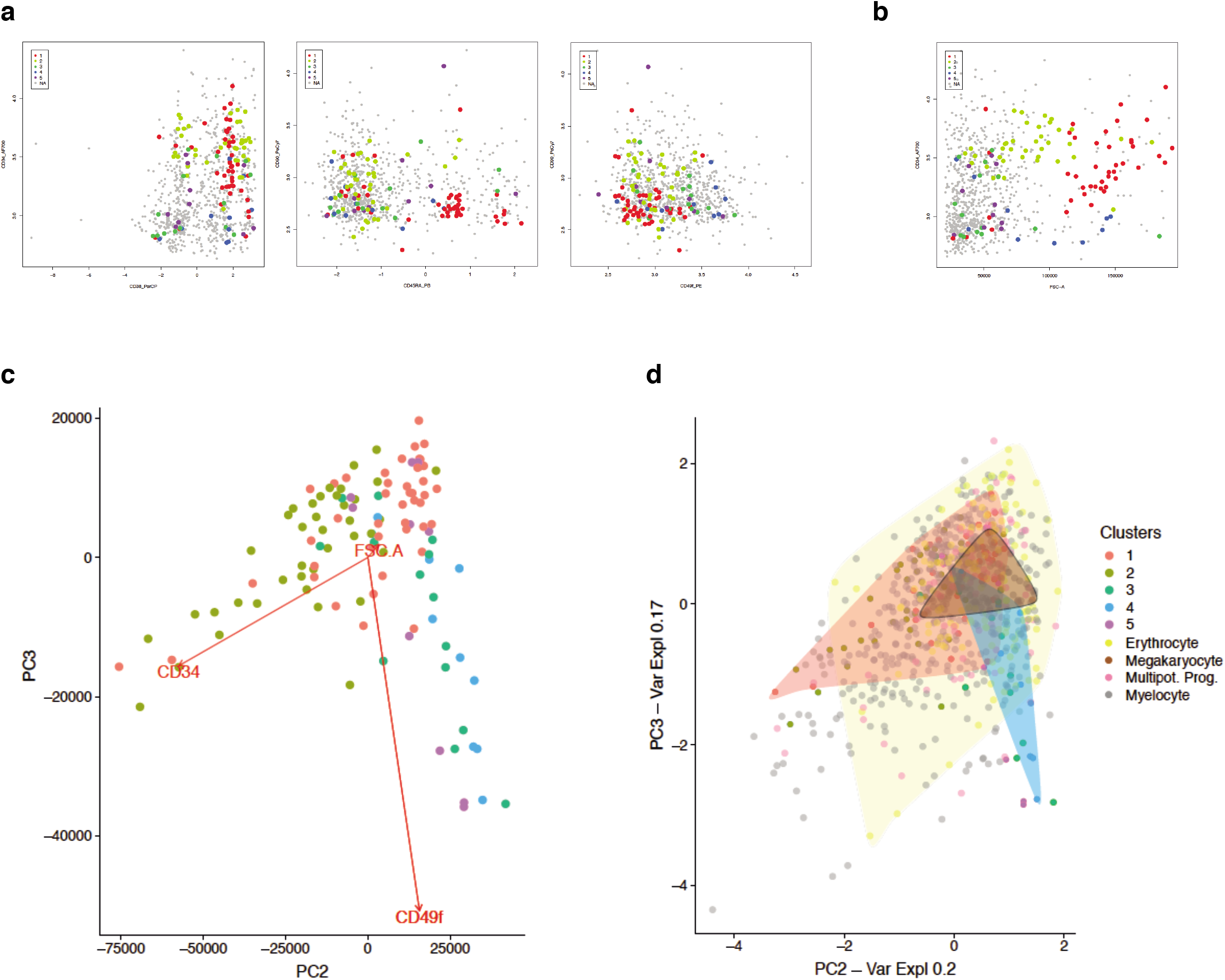
Prospective identification of HSC clusters. **a.** Retrospective analysis of FACS sorting strategy for each individual HSC sorted. Left panel: CD34 vs CD38; Middle panel: CD90 vs CD45RA; Right panel: CD90 vs CD49f. Individual surface marker expression was normalised. Cluster 1: red, cluster 2: yellow, cluster 3: green, cluster 4: blue, cluster 5: magenta, cells filtered due to poor quality: grey; **b.** Retrospective analysis of FACS index data: FSC-A vs CD34 for each individual HSC sorted. Identifies: cluster 1 (red): FSC-A^Hi^ CD34^Hi^, Cluster 4 (blue): FSC-A^Hi^ CD34^Lo^, cluster 2 (yellow): FSC-A^Lo^ CD34^Hi^. Individual surface marker expression was normalised. Cluster 1: red, cluster 2: yellow, cluster 3: green, cluster 4: blue, cluster 5: magenta, cells filtered due to poor quality: grey; **c.** Principal component analysis (PC2/PC3) of FACS surface marker expression for each individual HSC sorted including: FSC-A, SSC-A, Lin, CD34, CD38, CD45RA, CD90, CD49f. Vector loading for FSC-A, CD34 and CD49f are shown. cluster 1: red, cluster 2: olive, cluster 3: green, cluster 4: blue, cluster 5: magenta; **d.** Projection of FACS surface marker expression of HSCs characterized by differentiation assay and their lineage output onto Fig. 4c. Differentiation assay outputs: erythrocyte: yellow, MK: violet, multipotent progenitor: pink, myelocyte: grey. Cluster 1: red, cluster 2: olive, cluster 3: green, cluster 4: blue, cluster 5: magenta. Shaded areas: red: cluster 1, blue: cluster 4, yellow: erythrocyte output, violet: MK.

To characterize the lineage differentiation potential of Clusters 1 and 4, we compared our HSC index FACS profiles with those of Belluschi et al.^36^, who previously determined the lineage potential of index sorted single HSCs from human cord blood using an optimized single cell colony-forming cell assay, which supported myeloid/erythroid and MK lineage differentiation. Using an identical flow sorting protocol we could project our bone marrow derived HCS onto the PCA space of Belluschi *et al.* While only a small percentage of MK colonies containing MKs were observed after differentiation, when overlaid onto index data for HSCs in both studies, cells from HSC clusters 1 and 4 shared cell surface marker expression with HSCs forming MK colonies *in vitro* (Fig. 3d).

### Single cell and low input RNA sequencing of bone marrow MKs

To further characterize the nature of the development of bone marrow residing MKs we sequenced 188 pools of 20-100 MKs and 1106 single MKs. The MKs were isolated from fresh bone marrow harvested from 20 individuals undergoing sternotomy for heart valve replacement or coronary artery bypass grafting. The cell surface markers CD41 (GPIIb) and CD42 (GPIb) and the uptake of the DNA stain Hoechst 33342 was used to identify MKs of varying ploidy states (Fig. 4a).

**Figure 4.**
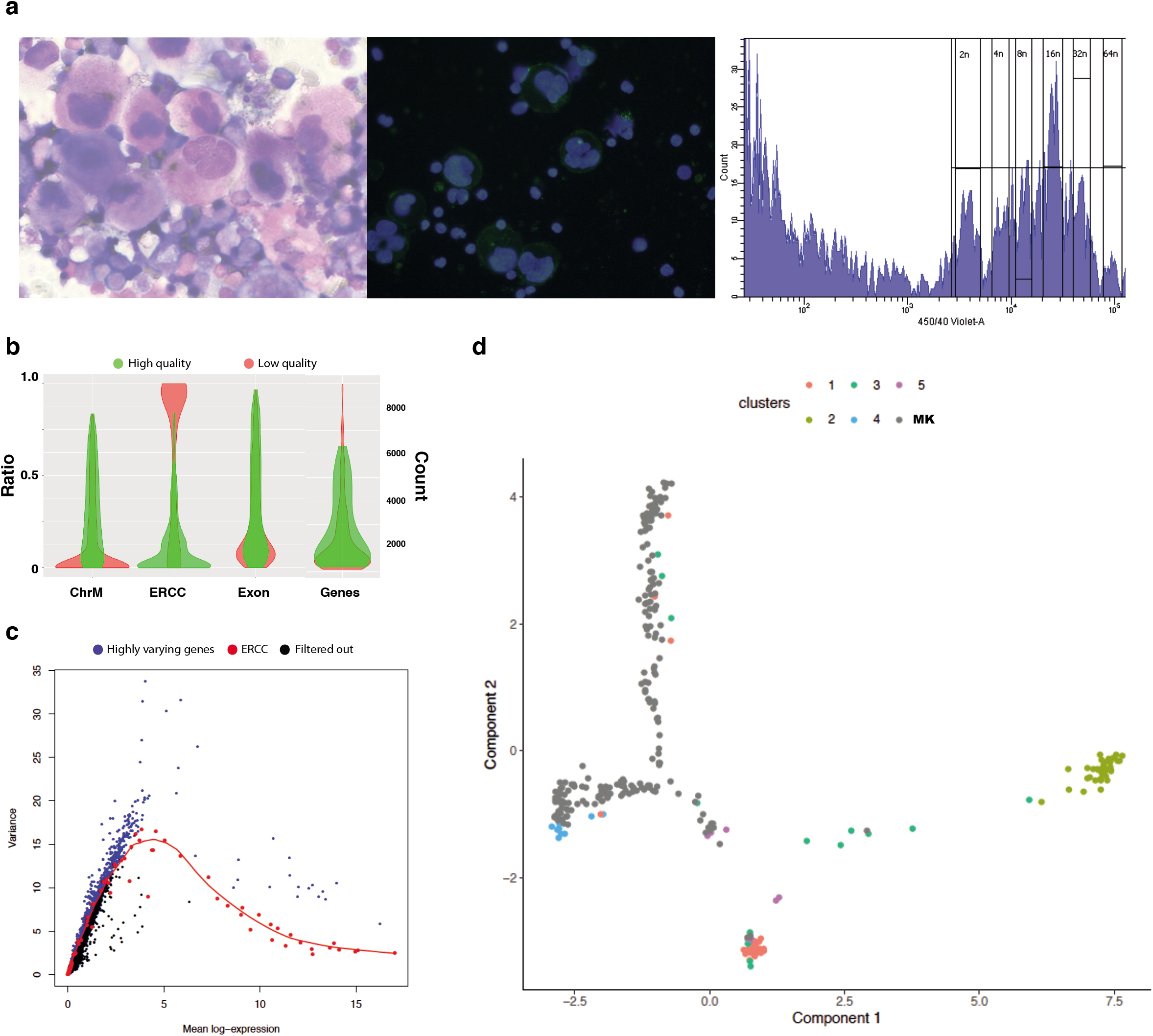
Single cell and low input RNA sequencing of bone marrow MKs. **a.** Human bone marrow MK phenotype. Left panel: Cytocentrifugation of enriched MK population stained with Roberts stain, 40x objective; Middle panel: Immunofluorescent staining of enriched megakaryocyte population with DAPI DNA stain (blue) and CD41 surface stain (green); Right panel: Fluorescence activated cell sorting for primary megakaryocytes from whole human bone marrow. Ploidy plot shown detecting levels of Hoechst 33342 staining showing typical ploidy distribution for human bone marrow MKs (cells shown are CD41a+, CD42a+); **b.** Single MK cell filtering. Performed using a 5-round training scheme of random forest models^37^ trained cDNA from 20 cell MK pools that were isolated using an identical sorting and sequencing protocol. High and low quality cells shown by mitochondrial mapping (ChrM), ERCC ratio (ERCC), exonic ratio (Exon), number of called genes (Genes). Green denotes high quality, red denotes low quality. Out of 1106 cells, 282 were taken forward for further analysis based on high quality; **c.** Gene filtering. Variance of normalized log-expression values for each gene in the HSC dataset, plotted against the mean log-expression. Variance estimates for ERCC spike-in transcripts and curve fit are highlighted in red. Blue dots represent highly varying genes (n=2000); **d.** Ordering differentiation trajectories between HSC cell clusters using Monocle 2 with the addition of the MK 20-100 cell pools. HSC clusters and MKs are differentiated by color.

Filtering of low quality single cells was performed with a 5-round training scheme of random forest models^37^ trained on the 20 cell MK pools that were isolated using the same methods. Using this model 282 single cells were taken forward for downstream analysis (Fig. 4b). ERCC spike-ins were again used for gene-wise normalization and filtering of highly variable genes where 2000 were taken forward for further analysis (Fig. 4c).

To investigate where the mature MKs transcriptionally fell in terms of differentiation we added the MK pools to the HSC trajectory analysis above using Monocle2^34^. The addition of mature MK transcriptional signatures to the HSC developmental trajectories showed that these are most similar to and branch from HSC clusters HSC clusters 1 and 4 with the MKs forming their own trajectory within the PCA (Fig. 4d).

### Development and maturation of bone marrow residing MKs

To investigate the changes in gene expression with increasing ploidy, we sequenced 32 pools of 20-50 MKs and 1106 single MKs with different ploidy (2N to 32N). The MKs were isolated from fresh bone marrow harvested from 10 individuals undergoing sternotomy for heart valve replacement (Supplementary Table S8).

In the 100 highest expressed genes from MK pools we found enrichment for mitochondrial and cell metabolism genes such as *MT-RNR1, MT-RNR2, CYTB* and *COX-1* (Supplementary Table S9) as previously described in platelet transcriptome studies^38^. Amongst these genes we observed a bimodal distribution in expression. 4N and 8N MKs shared expression of a number of genes (n=15) while there was a separate group of genes (n=40) mostly expressed in 16N and 32N cells, with different patterns of GO terms enrichment (Supplementary Table S10-S15). The genes characterizing the lower ploidy group were mostly involved in platelet function whereas, with further endomitotic replication there was a progressive enrichment in genes associated with protein processing and translation (Fig. 5a).

**Figure 5.**
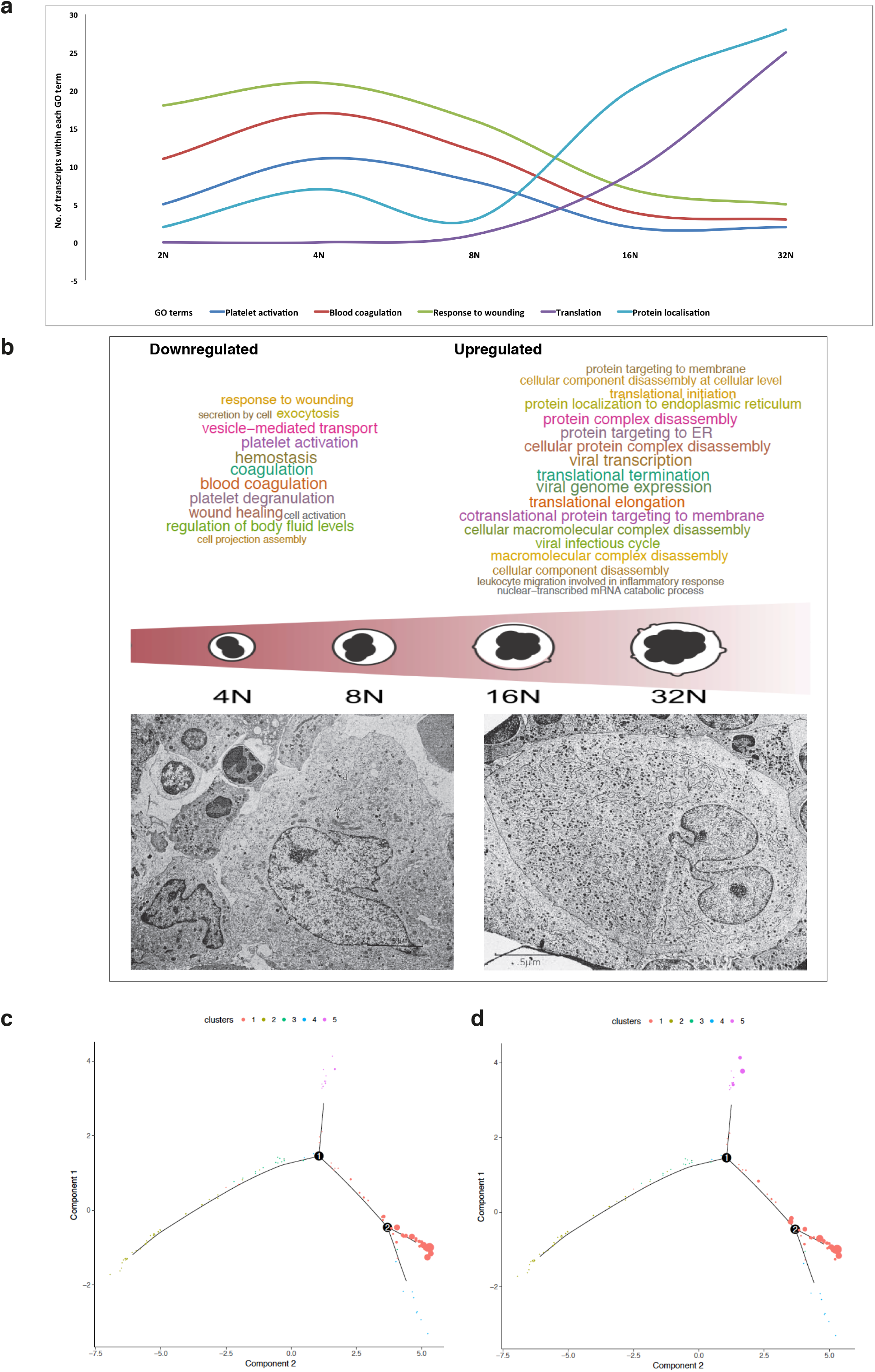
Development and maturation of bone marrow residing MKs. **a.** Number of genes expressed related to specific GO terms at each ploidy level, data shown based on 100 most abundantly expressed genes. GO terms: platelet activation, blood coagulation, response to wounding, translation, protein localization. The full GO analysis may be found in Supplementary Table S10-S14; **b.** Two distinct transcriptional states in MK differentiation. Upper panel: Over-represented GO terms in downregulated and upregulated genes with increasing ploidy, data shown is based on differential expression analysis between 32N and 4N MK 20 cell pools. The word cloud is a visual representation of the over-represented GO biological processes (size: indicator of significance). Full GO analysis may be found in Supplementary Tables S20,S21; Lower left panel: Electron microscopy image of a human stage 2 basophilic MK (2N-4N). Platelet specific granules (d), mitochondria (m), free ribosomes (r). Lower right panel: Electron microscopy image of a human stage 3 granular MK (8N-32N). Many platelet specific granules, mitochondria, rough endoplasmic reticulum. Black line or dotted black line of DMS is partitioning the MK cytoplasm into platelet territories. (Lower left and right panels reused from the Proceedings of the Japan Academy, Ser. B^39^ and the original figures were reproduced from Fig. 1.51, Fig. 1.52 respectively in Kosaki, G. and Fujimoto, T. (1979) Morphology of blood platelets, megakaryocytes differentiation and platelet release (in Nippon-Ketsuekigaku-Zensho (New Edition), vol. 11. Maruzen, Tokyo with permission from the publisher); **c.** Single cell HSC differentiation trajectories using Monocle 2 showing expression of MK ploidy signature: genes that are downregulated with increasing ploidy. Individual HSCs are connected by a minimum spanning tree with branch points (thin lines), representing the differentiation trajectory, clusters are differentiated by color. Expression of gene signature is indicated by the size of each cell; **d.** Single cell HSC differentiation trajectories using Monocle 2 showing expression of MK myocardial infarction signature. Individual HSCs are connected by a minimum spanning tree with branch points (thin lines), representing the differentiation trajectory, clusters are differentiated by color. Expression of gene signature is indicated by the size of each cell.

We performed differential gene expression (DGE) analysis with increasing ploidy using transcriptomes from MK pools as well as single cells for validation (Supplementary Tables S16-S19). We detected 944 DGE between 32N and 4N MK pools. 373 features were upregulated with increasing ploidy [False discovery rate (FDR)<0.05; with 181 features with an FDR <0.0001] and 571 features were downregulated (FDR<0.05, 179 with an FDR <0.0001). GO term enrichment analysis of the transcript biotypes revealed an enrichment for platelet degranulation, coagulation, hemostasis, wound healing and vesicle mediated transport in the lower ploidy MK transcriptome. Instead with increasing maturation and polyploidization these biotypes were downregulated in favor of terms related to translational initiation, elongation and termination, protein localization and cellular protein complex disassembly (Fig. 5b and Supplementary Tables S20 and S21). The same results were found in the single cell differential gene expression/GO analysis (Supplementary Tables S22 and S23).

Collectively, the MK transcriptional landscape changes with maturation from a low ploidy state, with the expression of large numbers of genes related to platelet function, to a high ploidy state, where there is an enrichment of genes involved in translation and protein packaging arguably in preparation for platelet release. This contemporary model of MK maturation is in direct contrast with the current understanding based on *in vitro* derived MKs, in which accumulating ploidy is transcriptionally associated with an upregulation of genes involved with platelet, coagulation and hemostatic pathways^24^. Our transcriptome data is, however, in keeping with observed changes in MK ultrastructure during *in vivo* maturation. While stage 2 MKs (low ploidy) have free ribosomes, smooth endoplasmic reticulum and few granules, stage 3 MKs (high ploidy) are characterized by rough endoplasmic reticulum, formed alpha and dense granules, increases in heterochromatin and smaller nucleoli indicating a reduction of transcription^39–41^ (Fig. 5b).

Using the expression profiles from increasing MK ploidy states (2N-32N), we modeled the trajectory from low ploidy to high ploidy, and were able to identify genetic programs turning on or off with the increasing of ploidy. This transcriptional signature was then plotted onto the HSC differentiation trajectories (Fig. 5c). Genes that are switched off at high ploidy are more active in cluster 1 HSCs. This reflects strong MK priming in HSCs and shows that gene expression modules activated during MK differentiation after branching point 2 are then gradually switched off as the cells reach maturation, in favor of alternative gene expression programs. This finding was confirmed using both single cell and MK pool sequencing data (Supplementary Tables S22 and S23). Of note, genes whose expression increases with increasing ploidy did not map to any specific HSC cluster.

### MK interactions within the bone marrow niche

The transcriptional differences that accompany increasing MK ploidy, highlighted in this work, are likely to underpin the functional differences observed between bone marrow MKs and *in vitro* derived MKs, namely efficiency of polyploidization and platelet release, which are likely to be due to signaling from the bone marrow niche. To investigate extracellular signals from the bone marrow niche that might contribute to MK maturation *in vivo*, we annotated genes upregulated with accumulating ploidy in the MK pool sequencing dataset in terms of their subcellular localization using the Ensembl database^42^ (Table 1). Of the 373 upregulated genes, 78 contained transmembrane domains and most likely localized to the cell membrane. While these have a wide range of functions, there are a number of genes coding for transmembrane proteins known to be expressed on the platelet membrane including: the G-protein coupled receptor PTGER2, the tetraspanin CD63, the TNF alpha receptor TNFRSF1B and lysophosphatidic acid receptor PPAP2A. *PTGER2, TNFRSF1B, P2Y6* encoding for the pyrimidine nucleotide receptor P2Y6, genes encoding for cytokine receptors IL13RA1, IL15RA as well as the complement receptor CR1 are highly enriched with increasing MK ploidy (FDR<1E-8). Many of these genes (except CD63) have extremely low levels of expression in previously published datasets of *in vitro* cultured MKs^25^ (Supplementary Fig. 7). We hypothesize that a number of these transmembrane proteins may mediate extracellular signals that drive MK maturation, increase in ploidy and platelet release.

**Table 1.**
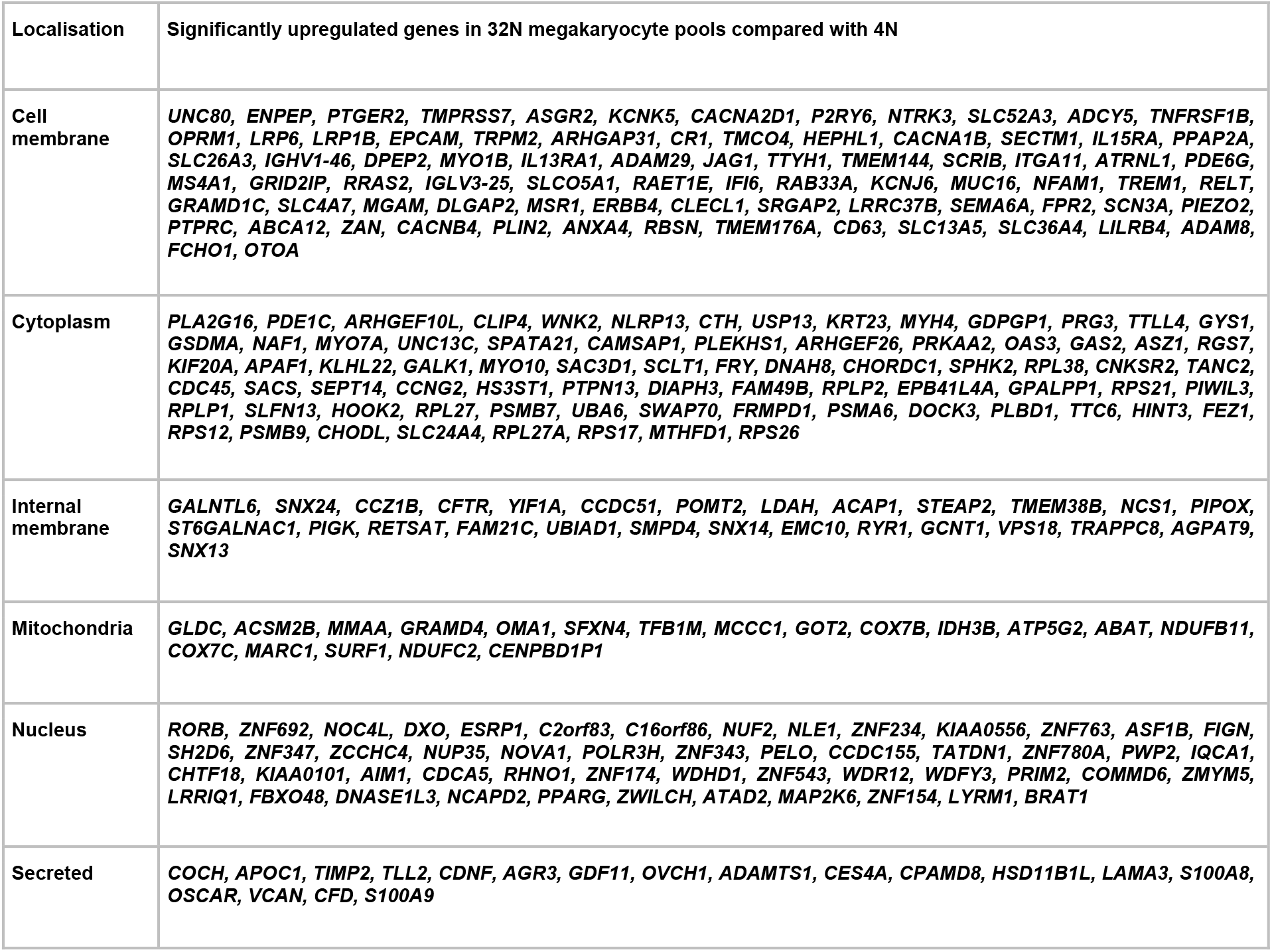
Transcripts upregulated with ploidy: Localisation within the cell. Using the Ensembl database the genes upregulated with ploidy were annotated in terms of their localization within the cell. 78 contained transmembrane domains and localized to the cell membrane, 77 were annotated as encoding cytoplasmic proteins, 57 as nuclear proteins, 27 as internal membrane proteins, 18 as mitochondrial proteins, 18 and secreted proteins and for 104 transcripts there was no localization information available.

### MK signature in myocardial infarction

Myocardial infarction has been suggested as a model for stress or accelerated thrombopoiesis accompanied by elevated mean platelet volume and reticulated platelet count^29^. We compared the transcriptomes of MK obtained from patients with severe coronary disease and recent myocardial infarction in the last 6 months (individuals undergoing sternotomy for coronary artery bypass grafting) with those from a control group (individuals undergoing sternotomy for heart valve replacement). We sequenced a total of 156 pools of 50-100 MKs (irrespectively of ploidy); 101 pools from 7 individuals with severe coronary disease and recent myocardial infarction and 55 pools from 8 controls (Supplementary table S24).

DGE analysis revealed 139 upregulated (FDR<0.05; with 21 features with an FDR <0.00010 and 679 downregulated (FDR<0.05; with 62 features with an FDR <0.0001) genes in MKs from patients with severe coronary disease and recent myocardial infarction. A number of upregulated genes were directly related to platelet activation and proteins released by the alpha granule including *PPBP*, *THBS1* and *RAP1B* as well as the glutamate receptor GRIA1.

We derived MK gene signatures associated with myocardial infarction and coronary disease and visualized these signatures onto the HSC differentiation trajectory plot (Fig. 5d). Genes that are modulated with coronary disease were active in Cluster 1 HSCs. None of the MK genes regulated by disease aligned to Cluster 4 HSCs.

## Discussion

Here we used single cell RNA sequencing to chart the transcriptional journey of human bone marrow MKs, from HSC early lineage priming, showing the presence of 2 separate MK primed clusters along the differentiation trajectory, to the later stages of MK maturation and polyploidization.

We identified 5 transcriptionally distinct HSC subsets in human bone marrow using three independent clustering methods. Two of these subsets (cluster 1 and cluster 4) displayed transcriptional priming towards the MK lineage and indeed when overlaid onto data from single cell assays^36^, they showed a functional MK lineage bias. Cells in these two clusters expressed genes that have been shown to be highly expressed in the MK-platelet lineage in previous studies^22,25,35,38^. The high proportion of HSCs displaying priming to the MK lineage is in keeping with recent evidence indicating that this is the first lineage bifurcation in the multipotent HSC compartment^4^ and the role of MKs in HSC regulation within the stem cell niche^15,18,43^. Plotting these cells in pseudotime differentiation trajectories showed that cluster 1 and cluster 4 in fact branch from the same trajectory. Comparison with previous datasets^25,35^ demonstrated that, while cells in cluster 1 were enriched for genes involved in HSC function, megakaryopoiesis and genes commonly expressed between MK and MEP populations, cells in cluster 4, in contrast, were enriched for genes specific to MKs with a number of marker genes for this cluster directly related to platelet function. We have therefore delineated single cell heterogeneity within the bone marrow HSC compartment highlighting 2 subpopulations that are potentially unilineage MK.

Our work presents the first interrogation of the transcriptional landscape of human bone marrow MKs. The knowledge of the MK transcriptome until now being based largely on *in vitro* derived MKs from CD34+ cells^21–25^. This analysis was made possible by the advances in single cell and low input sequencing techniques. We recreated a differentiation trajectory combining the HSCs and MKs transcriptomic data together and we have shown that human bone marrow MKs branch directly from phenotypic HSCs, specifically cells in clusters 1 and 4 supporting the notion of transcriptional priming of these clusters towards the MK lineage.

We charted transcriptional changes in human bone marrow MKs through different stages of polyploidization. Microarray data of increasing ploidy in *in vitro* derived MKs has previously shown an overall pattern of upregulation of genes involved with platelet, coagulation and hemostatic pathways and a downregulation of cell cycle associated genes^24^. Our results challenge this view. Our data provide a model of MK development and polyploidization whereby two distinct transcriptional states exist for low and high ploidy MKs and the cells switch from one to the other with successive endomitotic replication. With increasing ploidy we observed a downregulation of genetic programs related to platelet functionality, although these genes are still switched on and remain expressed within the higher ploidy MKs. Conversely, with increasing ploidy level, we observed a marked upregulation of genes encoding ribosomal subunits, those associated with protein translation and protein localization. These findings indicate that at high ploidy the energy is redirected to mRNA translation and appropriate localization of the newly generated proteins into alpha granules, dense granules and other vesicles, readying the cell for thrombopoiesis and platelet activation. Bone marrow MKs high energy requirement, particularly those of higher ploidy, is the likely explanation for the high levels of mitochondrial and other genes involved in cellular metabolism since the production of ribosomes consumes large amounts of energy^44^. Our data are in keeping with electron microscopy studies of MK ultrastructure at different points of maturation^39–41^.

When the MK ploidy signature was plotted on to HSC differentiation trajectories, it overlapped with gene expression in cluster 1 HSCs only. This signature was modeled on genes that were downregulated with increasing ploidy indicating that genes modulated in cluster 1 HSCs are important in MK lineage priming and early MK development in steady state thrombopoiesis.

Identifying genes upregulated with increasing ploidy also has implications for improving our understanding of MK platelet release. Currently, the number of platelets produced for each *in vitro* generated MK is ~1000-fold lower than *in vivo*, which has severe implications for platelet production for clinical transfusion from iPSC derived MKs^45^. This observation is likely a direct result of the differences between MKs produced in culture and MKs in the bone marrow niche in terms of ploidy level, cytoplasmic maturation or extracellular signals. A number of genes upregulated with increasing ploidy were identified to encode transmembrane receptors *(PTGER2, TNFRSF1B, IL13RA1* and *PPAP2A)*, which upon binding to their ligands, might modulate functional effects such as MK chemotactic migration from the osteoclastic niche to the vascular sinusoidal space or indeed initiate platelet production. These may represent important novel drivers of MK maturation and platelet release. PTGER2, a G-protein coupled receptor known to be present on the platelet membrane, is an important regulator of HSC expansion and function in the bone marrow niche^46^ and has been shown to specifically promote megakaryocyte lineage recovery after radiation injury in mouse^47^. Binding of TNFRSF1B, also present on the platelet membrane, suppresses apoptosis and has been shown to induce megakaryopoiesis in hematopoietic progenitors which could in part account for increased platelet mass in inflammation^10^. IL13RA1 mediates the effects of IL13 which significantly increase MK colony formation *in vitro*^48^ and in murine models^49^. PPAP2A is an integral membrane glycoprotein that degrades lysophosphatidic acid (LPA) by dephosphorylation. LPA has been shown to inhibit megakaryopoiesis *in vitro*^50^. Furthermore, an LPA gradient has been proposed between the osteoblastic niche and vascular sinusoid regulating MK localization and maturation^51^. Future work on these transmembrane receptors and testing appropriate ligands to investigate their role in MK maturation would be imperative in identification of novel drivers of MK maturation.

We also compared MK transcriptional signatures in myocardial infarction as a model for stress thrombopoiesis with controls. Our data showed that a number of genes upregulated in MKs from individuals with severe coronary disease and myocardial infarction compared to controls were related to platelet activation such as *PPBP* and *THBS1* and *RAP1B* that encodes for a protein stimulated by collagen binding to and plays a critical role in modulating the affinity state of GP2b/3a^52^. RAP1B has also been identified as a potential therapeutic target in myocardial infarction^53^. There is now compelling evidence for enhanced megakaryopoiesis as an important pathogenic driver in atherosclerosis and myocardial infarction^54,55^. The clinical trial CANTOS^56^ demonstrated an improvement in cardiovascular outcome in patients with myocardial infarction with Canakinumab, a monoclonal antibody targeting IL1B, a known driver of megakaryopoiesis both *in vitro* and *in vivo*^57^. We found that the gene encoding the AMPA glutamate receptor, *GRIA1*, was upregulated in MKs in individuals with myocardial infarction. As glutamate serum levels are increased in thrombosis^58^ and interruption of glutamate binding to the NMDA glutamate receptor in megakaryocyte cell lines has impaired megakaryopoiesis^59^, this observation raises the possibility of a positive feedback mechanism of glutamate signaling leading to increased platelet production perpetuating a prothrombotic state. Therefore our data also supports a pathological role of stress thrombopoiesis in acute coronary thrombosis.

Finally, we plotted the gene signature obtained from MK myocardial infarction onto the HSC differentiation trajectories and we showed that it overlapped with the genes active in cluster 1 HSCs, in keeping with the MK ploidy signatures. This suggested that the MK lineage priming used in steady state thrombopoiesis potentially also occurs in stress thrombopoiesis.

Taken together our work describes the transcriptional heterogeneity within the human bone marrow HSC population with 2 subpopulations exhibiting MK lineage priming and direct branching of MK differentiation from these HSC subpopulations. Furthermore, it charts the transcriptional journey from MK lineage commitment through to the mature polyploid MK and apply this to the setting of myocardial infarction. Ultimately better understanding of MK lineage commitment, differentiation from the HSC, maturation and platelet release are key to deciphering pathological modulation leading to thrombosis and bleeding disorders.

## Methods

### Human samples

Pseudonomised bone marrow for HSC and MK sorting was obtained from individuals undergoing cardiac surgery at Barts Health NHS Trust, London, after informed consent and ethical approval from London - City & East, REC 13/LO/1760 (BAMI Platelet Sub-s. Individuals recruited fell into two separate categories: 1. elective open heart valve replacement with no evidence of coronary artery disease on coronary angiography; 2. coronary artery bypass grafting in the context of recent myocardial infarction (within 6 months). Myocardial infarction was defined on the basis of clinical history, troponin rise and changes consistent with ischaemia on electrocardiography. Exclusion criteria comprised emergency surgery, presence or history of hematological malignancy, abnormal platelet count and haemoglobin levels <85g/l.

A sternal bone marrow scraping was taken directly following median sternotomy using a Volkmann’s spoon. The sample was collected into an EDTA Vacutainer tube containing 1.8mg/ml EDTA. 4mL of Dulbecco’s phosphate buffered saline (PBS, Sigma) containing 10% human serum albumin (HSA, Gemini Bio Products) was added and the whole volume was resuspended by pipetting 2-3 times. The sample was then put on metallic thermal beads (ThermoFisher Scientific) at a temperature between 0-4°C and transported to the University of Cambridge for further processing.

### Isolation of HSCs and MKs from fresh human bone marrow

Processing of clinical bone marrow took place 2-3 hours after harvest. The cellular content was flushed out of the bone marrow using PBS containing 1.2% HSA, 2mM EDTA (Sigma) and the red cells were lysed using ammonium chloride lysis.

For HSC isolation the cells were stained with the following antibody cocktail: PECy5 conjugated anti-lineage specific antibodies: CD2 (BD), CD3 (BD), CD10 (BD), CD11b (BD), CD11c (BD), CD19 (BD), CD20 (BD), CD56 (BD), biotinylated CD42b (Pab5, NHS Blood and Transplant, International Blood Group Reference Laboratory [IBGRL]), biotinylated GP6 (Pab5, NHS Blood and Transplant, International Blood Group Reference Laboratory [IBGRL]) used in combination with PECy5 conjugated streptavidin (Biolegend). Alexa Fluor 700 conjugated anti-CD34 (BD), PerCP-Cy5.5 conjugated anti-CD38 (BD), Pacific Blue conjugated anti-CD45RA (Invitrogen), PECy7 conjugated anti-CD90 (BD),PE conjugated anti-CD49f (BD). After staining cells were kept at 4°C before sorting using a FACS Aria Fusion flow sorter (BD). Single HSCs defined as Lineage-, CD34+, CD38-, CD45RA-, CD90+, CD49f+ cells were sorted by FACS directly into individual wells of a 96-well plate. Index sort data was collected for each single cell. To ensure that the cells that were sorted and studied were in fact HSCs with both multilineage engraftment potential, cells sorted within these gates were transplanted into sub lethally irradiated NSG (NOD scid gamma) mice and demonstrated both myeloid and lymphoid engraftment at 16 weeks.

For MK isolation the cells were stained for surface MK markers with mouse anti-human CD41a APC conjugated antibody (BD) and mouse anti-human CD42b PE conjugated antibody (BD) and for ploidy analysis with 1ug/ml Hoechst 33342 (Invitrogen). After incubation at 37°C for 30 minutes, the cells were kept at 4°C before sorting using a FACS Aria Fusion flow sorter (BD). Single cells and MK pools of 20 cells were sorted by FACS according to ploidy level using a 100uM nozzle directly into individual wells of a 96-well plate.

For both HSC and MK isolation cells were sorted into wells each containing 2.5ul lysis buffer/well: 950uL RLT Plus buffer (Qiagen) and 50ul 20U/ul SUPERase In RNase inhibitor (Ambion) and stored at −80°C until library preparation.

### RNA sequencing library preparation

cDNA synthesis and poly(A) enrichment was performed following the G&T-seq protocol^60^, a variation of the Smart-seq2 protocol. ERCC spike-in RNA (Ambion) was added to the lysis buffer in a dilution of 1:4,000,000. Streptavidin-coupled magnetic beads (Dynabeads, Life technologies) were conjugated to the biotinylated oligo-dT primer: 5’-biotin-triethyleneglycol-AAGCAGTGGTATCAAC GCAGAGTACT_30_VN-3’ where V is A/C/G and N is any base (IDT) according to manufacturer’s instructions. Separation of DNA and RNA was performed using a Biomek FXP Laboratory Automation Workstation (Beckman Coulter). 10ul conjugated beads were added to the cell lysate and incubated for 20minutes at room temperature with continuous mixing. The mRNA bound to streptavidin beads was then magnetized to the side of the well and the genomic DNA containing supernatant was transferred to a different plate. The beads were further washed 4 times at room temperature with a wash buffer consisting of 50 mM Tris-HCl pH 8.3, 75 mM KCl, 3 mM MgCl2, 10 mM DTT (Life technologies), 0.5% Tween-20, 0.1x 20U/ul SUPERase In RNase inhibitor.

After this either the Superscript II (MKs) or the Smartscribe (HSCs) reverse transcription (RT) mix (10ul) was added to the beads. Common to both mixes were: 0.25ul 20U/ul SUPERase In RNase inhibitor, 2ul 5M betaine (Sigma), 0.06ul 1M MgCl_2_ (Life technologies), 1ul 10mM dNTP mix (Thermo Scientific) and 1ul 100mM Template-Switching Oligo: 5’-AAGCAGTGGTAT CAACGCAGAGTACrGrG+G-3’ where rG is a ribo-guanosine and +G is a locked nucleic modified guanosine (Exiqon). The Superscript II RT mix also contained: 0.5ul 200U/ul Superscript II reverse transcriptase (Life technologies), 2ul 5x Superscript II First-Strand Buffer (Life technologies), 0.5ul 100mM DTT (Life technologies) and 3.59ul nuclease-free water (Life Technologies). The Smartscribe RT mix also contained: 1ul Smartscribe reverse transcriptase (Clontech), 2ul 5x Smartscribe First-Strand Buffer (Clontech), 1ul 20nM DTT (Clontech) and 2.59ul nuclease-free water. cDNA first strand synthesis was performed by incubating at 42°C for 60 min, followed by 50°C for 30 min and 60°C for 10 min with mixing on a Thermomixer (Eppendorf).

Global amplification of the cDNA by PCR was then performed by adding PCR mastermix consisting of: 12.5ul 2x KAPA HiFi HotStart ReadyMix (KAPA Biosystems), 0.25ul 10mM Smart PCR primer: 5’-AAGCAGTGGTATCAACGCAGAGT-3’ (Biomers) and 2.25ul nuclease-free water to the reverse transcription reaction mixture. The PCR reaction consisted of heating to 98°C for 3 min, 22 PCR cycles (98°C/20 s, 67°C/15 s, 72°C/6 min) and incubation at 72°C for 5 min. A 1:1 Ampure XP (Beckman Coulter) PCR purification step was performed followed by assessment of size distribution using an Agilent high-sensitivity DNA chip (Agilent technologies) and expression of selected genes was also examined at this stage using Taqman real-time qPCR. Briefly, TaqMan Fast Advanced Master Mix (Applied Biosystems) and Taqman probe: Glyceraldehyde-3-phosphate dehydrogenase (GAPDH, Hs02758991_g1) (Applied Biosystems) were used to perform real-time PCR reactions over 40 cycles according to manufacturer’s instructions on the StepOnePlus system. At least 2 technical replicates for each sample were performed.

Illumina libraries were prepared from between 0.5 and 10ng of amplified cDNA using the Nextera XT DNA sample preparation protocol (Illumina) and a 1:1 Ampure XP purification step was performed. Library size distribution was checked on an Agilent high-sensitivity DNA chip and the concentration of the indexed library was determined using the KAPA library quantification kit (KAPA Biosystems) according to the manufacturer’s instructions on the StepOnePlus system (Applied Biosystems). 150 base pair (bp) paired-end sequencing was performed on the Illumina HiSeq 4000 instrument using TruSeq reagents (Illumina), according to manufacturer’s instructions.

### RNA sequencing analysis

Paired-end reads were trimmed of PCR and sequencing adapters <32bp using TrimGalore! (http://www.bioinformatics.babraham.ac.uk/projects/trim_galore/ 2014). The reads were mapped to the human reference genome (GRCh37) using STAR^61^. Quality metrics were assessed for each sample including: total reads, alignment rate, ERCC ratio, number of called genes, ratio of exonic reads, ratio of reads mapping to the mitochondrial genes. These metrics were illustrated using: http://servers.binf.ku.dk:8890/sinaplot/^62^.

### Single cell RNA sequencing analysis

For HSCs filtering of low quality cells was performed using similar SVM approach and rafures as described^30^ using HSC single cell cDNA positive for the house-keeping gene *GAPDH* as determined by real-time qPCR as target for classification. For MKs filtering of low quality cells was performed using the same features in a 5-round training of random forest models^37^, using output of previous model as input for next model in order to iteratively increase the sensitivity, when trained to classify the 20 cell MK pools that were isolated using an identical sorting and sequencing protocol, when all other cells acted negatives.

The most highly variable genes were filtered above technical noise using the Scater package^63^ using previously described methods^32^. ERCC spike-ins were used to model the trend in technical variability.

Unsupervised clustering of single cells was performed in a principal component analysis (PCA) space using correlation distances correlation in the PCA space. The robustness of the clusters was determined by the Silhouette index^64^. Using SC3^65^, cluster marker genes were identified based on their predictive value to separate each cluster from the rest. A p-value was then assigned to each gene using the Wilcoxon signed-rank test. Genes with P<0.001 were defined as marker genes.

Cells were ordered into differentiation trajectory using the Monocle 2 single-cell analysis toolset^66,34^.

In order to assess expression of known hematopoietic gene signatures within the single cells clusters, gene signatures for each cell type in the DMAP dataset^35^ were created using the LIMMA package^67^. Here a one vs all comparison was made for all cell types. Genes that were considered to be part of the gene signature log2 fold change of >1 and FDR <0.05. The expression of genes from each DMAP gene signature was then assessed in each individual single cell and this information was superimposed as size of dots on the Monocle 2 plots. Furthermore, expression of cluster marker genes within hematopoietic progenitor populations in the Blueprint dataset^25^ was assessed using the Blueprint tools portal: https://blueprint.haem.cam.ac.uk.

### Differential expression analysis

DESeq2^68^ was used to find genes that were differentially expressed using the Wald test, and size factors estimated by scran^69^. Differentially expressed genes were defined as FDR<0.05.

### Functional gene list analysis

Gene ontology (GO) analysis was performed and its visual representation generated using the online tool, Fidea^70^: http://circe.med.uniroma1.it/fidea. Here enrichment analysis is performed by statistically assessing whether a pathway or process is enriched in the specific gene list. This is achieved by the hypergeometric test with the resulting p-values corrected using the Benjamini and Yekutieli FDR method.

### Analysis of indexed FACS data

FACS data from previous study by Belluschi *et al.*^36^ were used in PCA analysis on scaled data (PCA_B), onto which new samples (BM_HCS) where projected by applying PCA_B scaling target and center, and the position was calculated as dot-product of scaled BM_HSC vector and PCA_B rotation. Cells from the previous study were then marked according to differentiated cell type, as determined by FACS, where cell type perimeter is shown via geom_encircle (https://CRAN.R-project.org/package=ggalt) using parameters s_shape=0.6 and expand=0.1.

## Data availability

All RNA-seq data have been deposited in the European Genome-phenome Archive under the accession number______. All relevant data are also available from the authors.

## Author contributions

F.A.C. designed and performed experiments and wrote the manuscript; F.O.B. performed bioinformatic analysis and wrote the manuscript. I.C.M supervised experiments and parts of the study. S.F., F.B., C.K., H.M. and K.D. carried out experiments. L.R.O. and N.H provided bioinformatic analysis and edited the manuscript. A.M, J.F.M, R.U supervised the clinical aspects of the study. T.V., W.H.O., E.L. and S.T. supervised parts of the study and edited the manuscript. M.F. designed and supervised experiments and wrote the manuscript. F.A.C. and F.O.B contributed equally to this work.

---

**Supplementary Figure 1.**
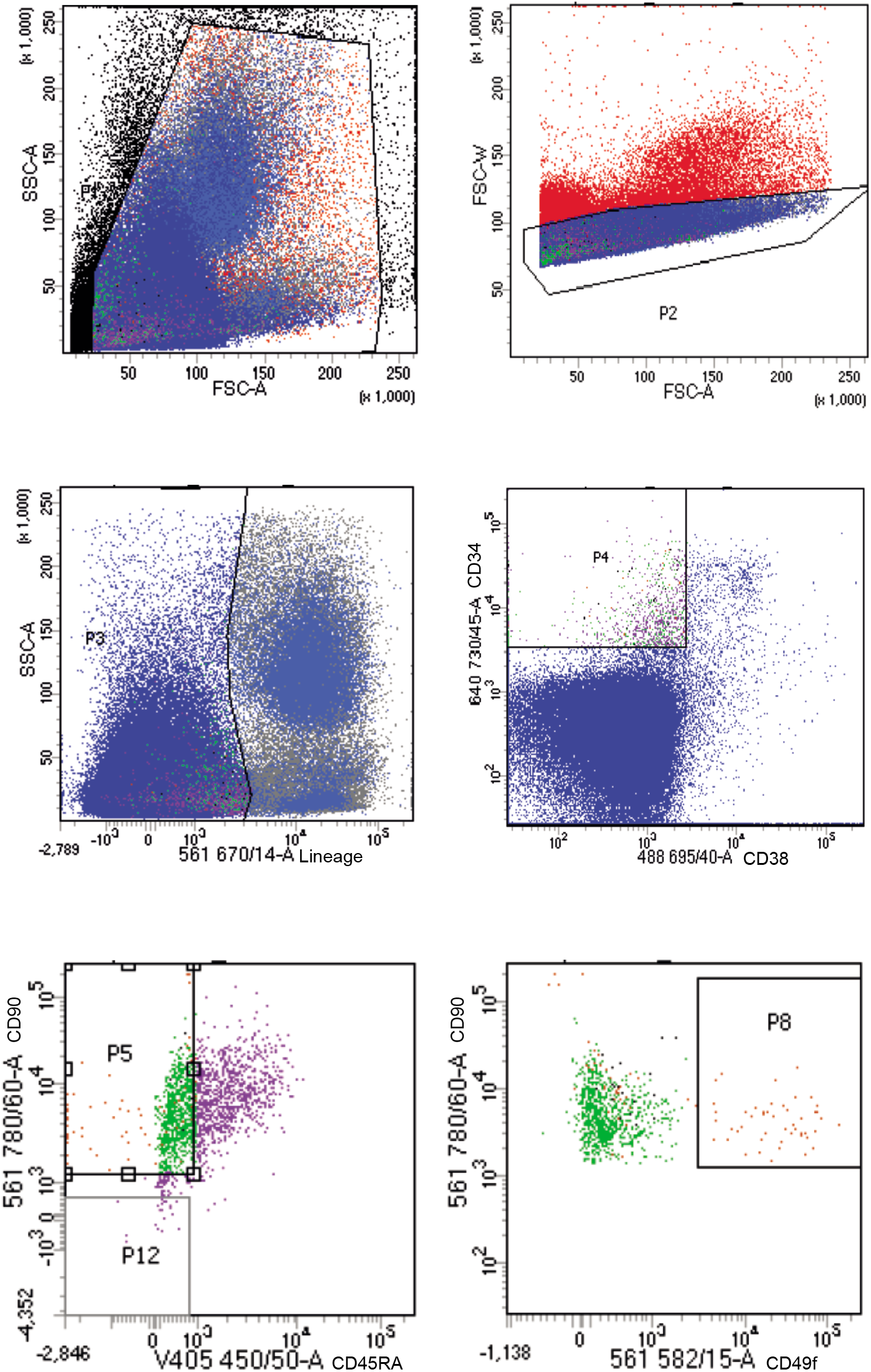

**Supplementary Figure 2.**
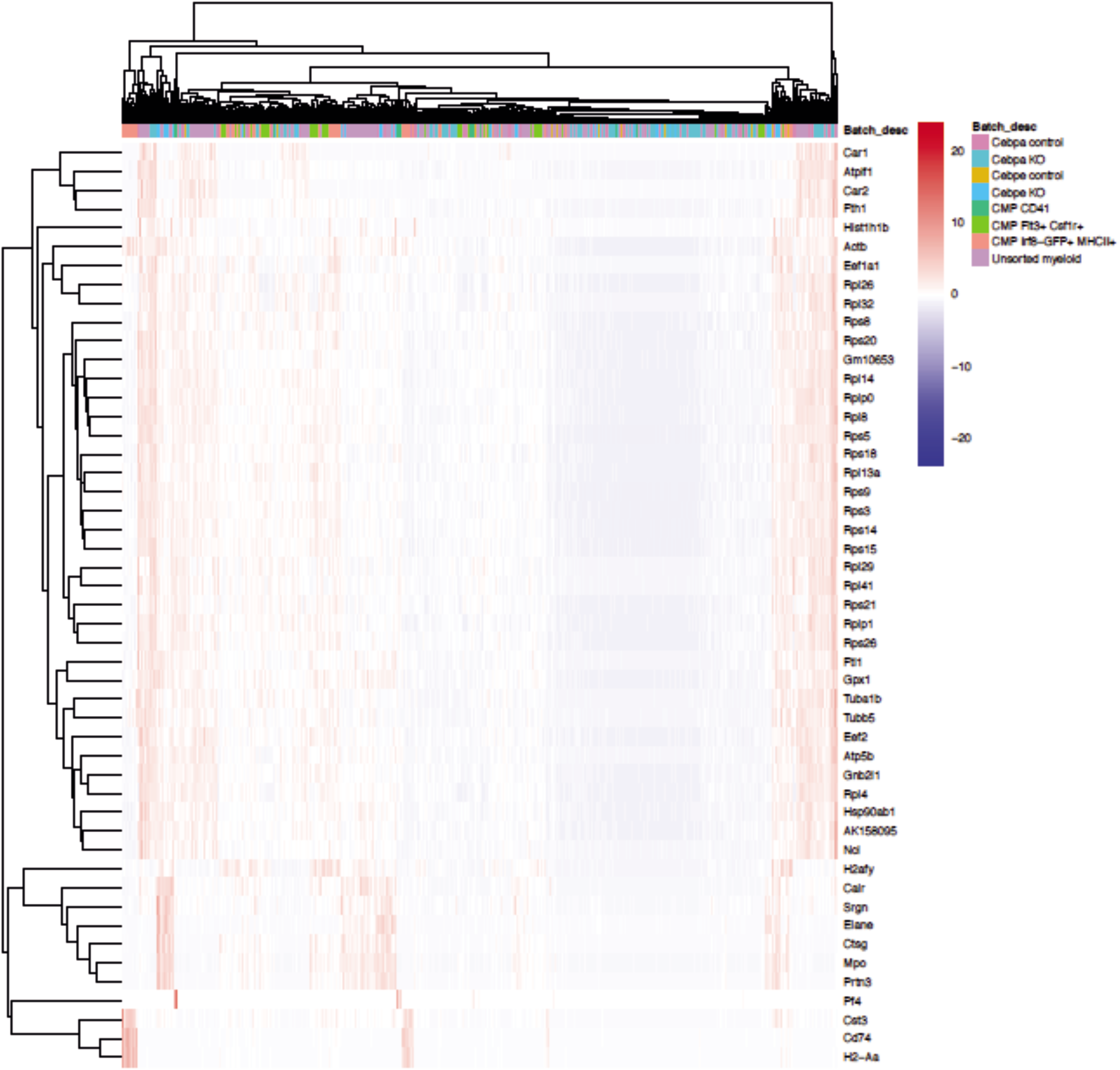

**Supplementary Figure 3.**
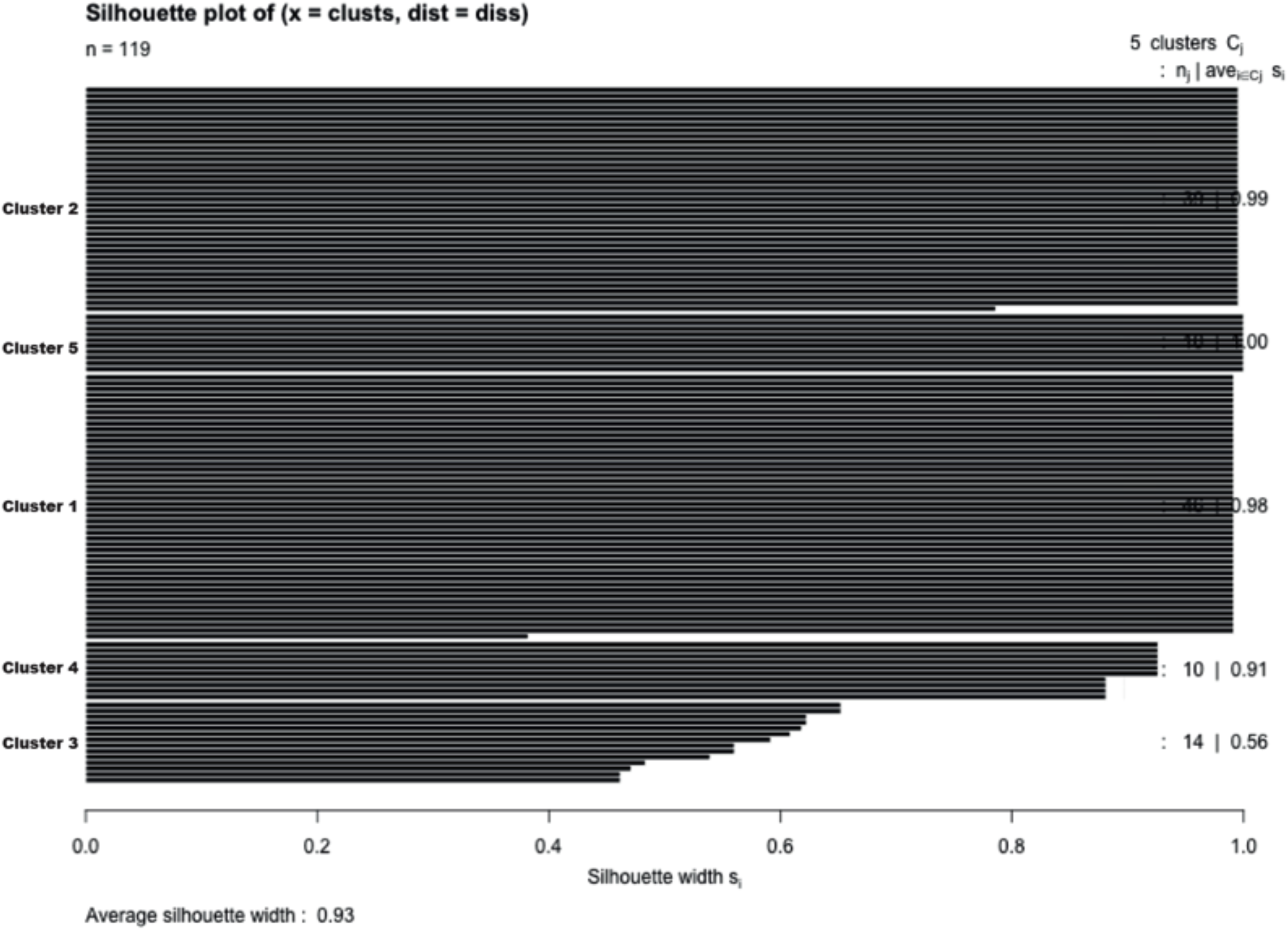

**Supplementary Figure 4.**
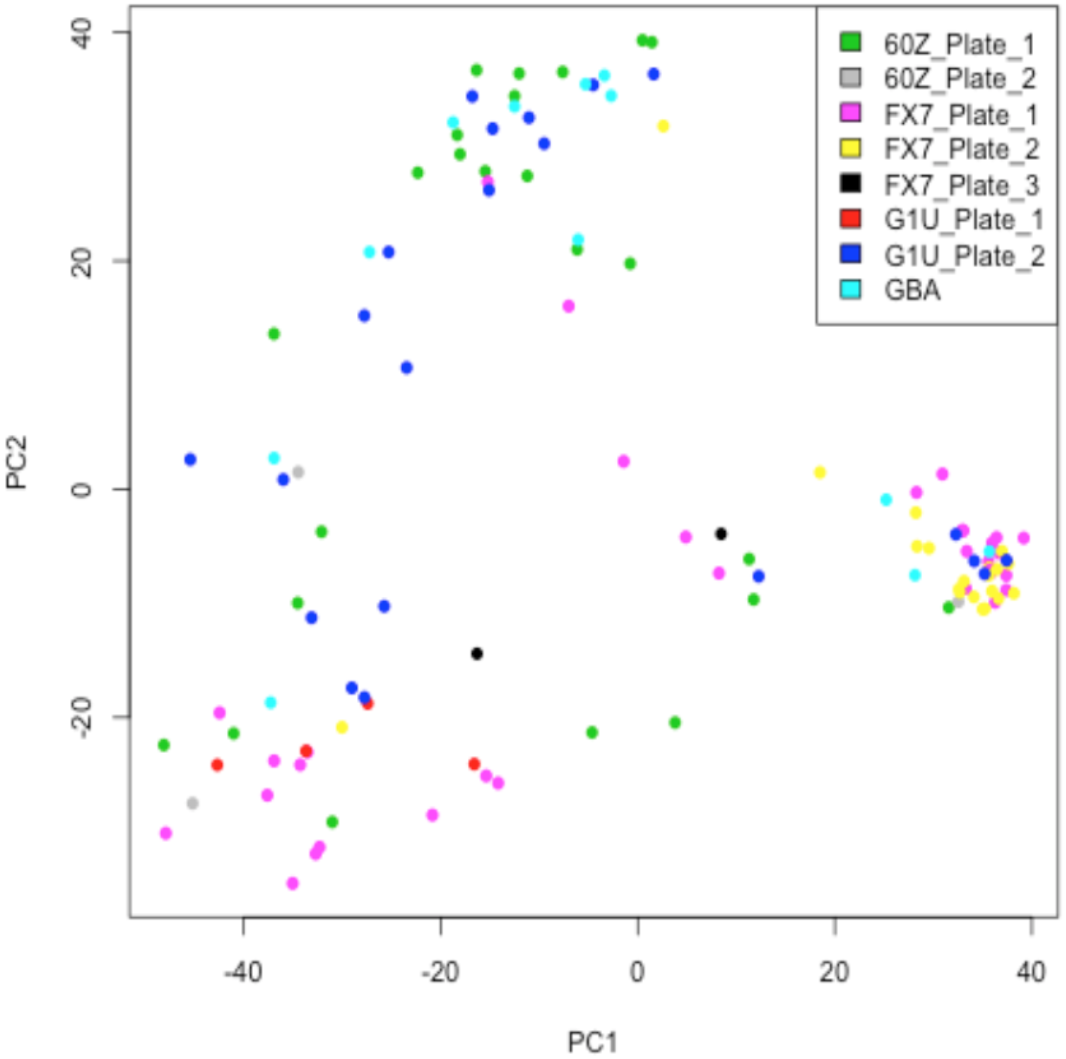

**Supplementary Figure 5.**
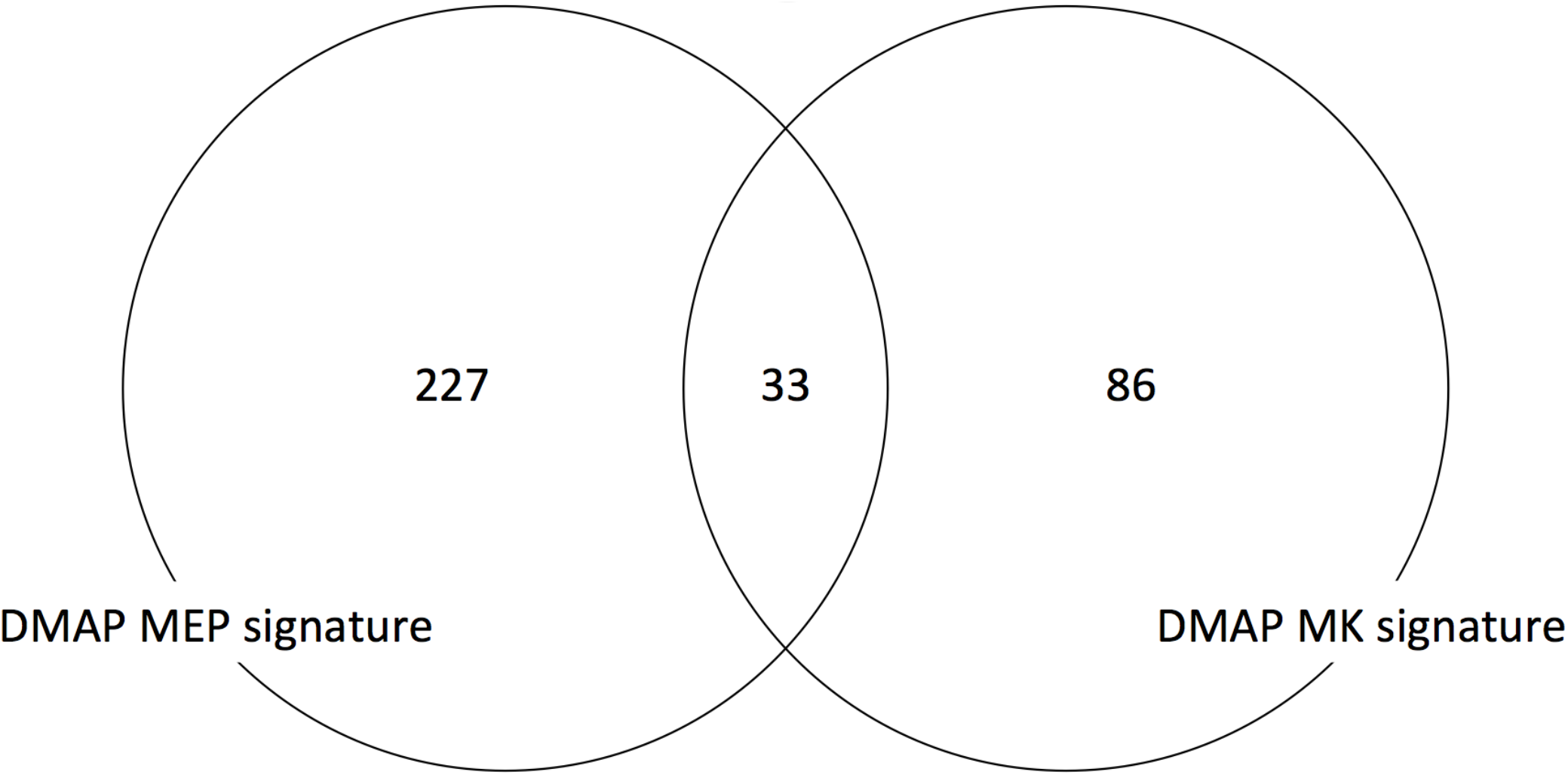

**Supplementary Figure 6.**
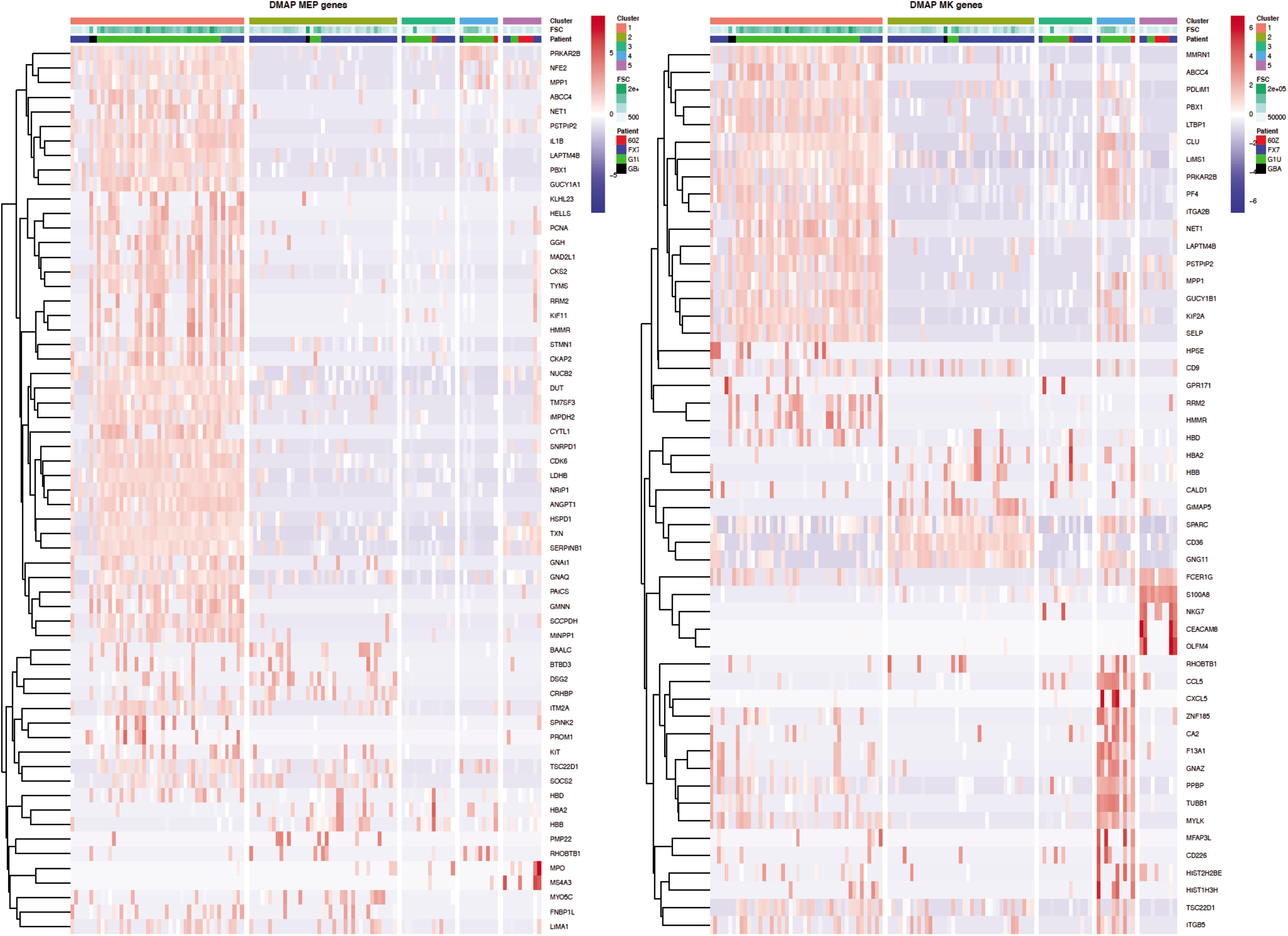

**Supplementary Figure 7.**
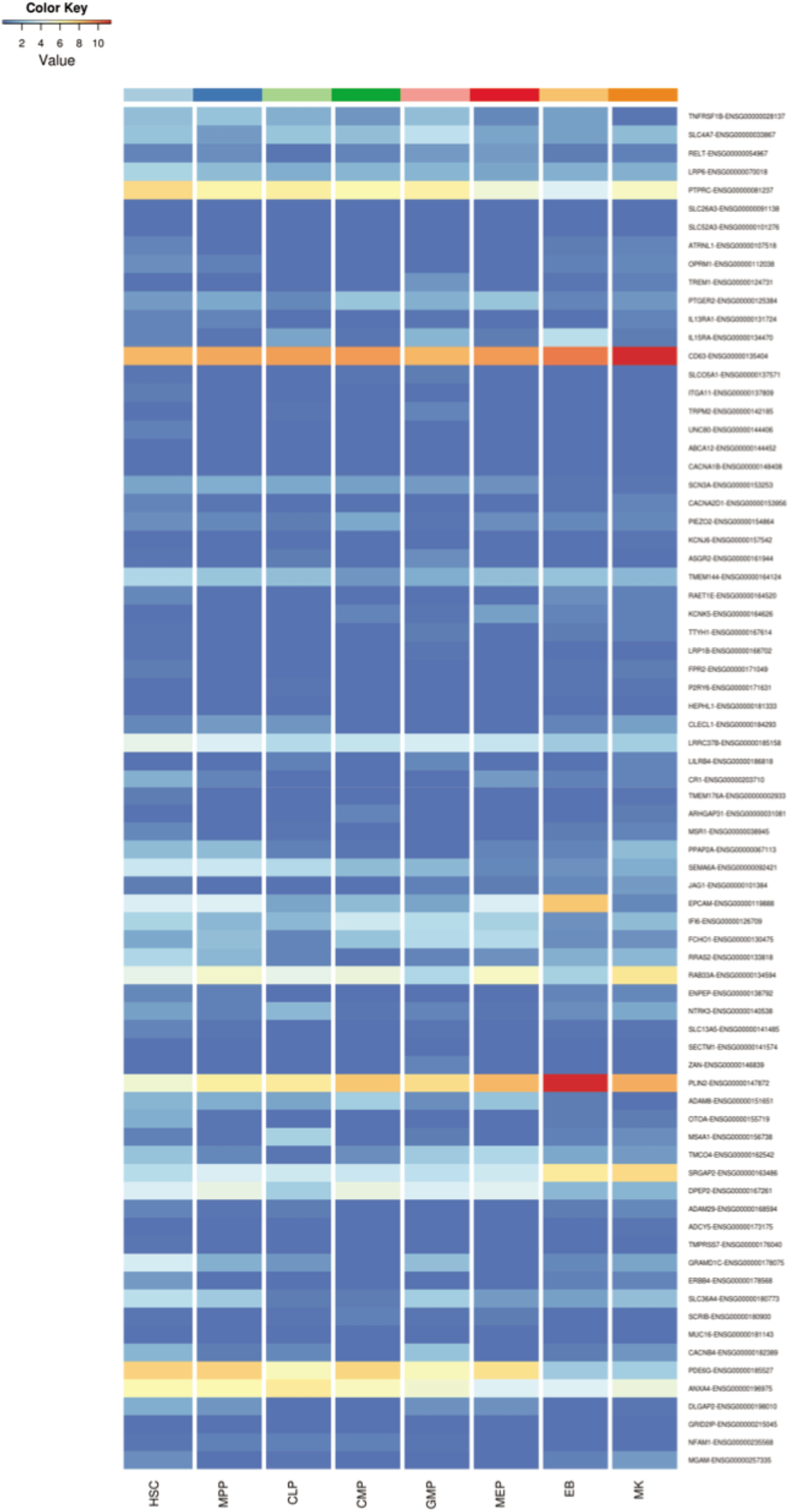

